# F2,6BP restores mitochondrial genome integrity in Huntington’s Disease

**DOI:** 10.1101/2024.11.04.621834

**Authors:** Anirban Chakraborty, Santi M. Mandal, Mikita Mankevich, Rajam S. Mani, Shandy Shahabi, Tapan Biswas, Manohar Kodavati, Sravan Gopalkrishnashetty Sreenivasmurthy, Nisha Tapryal, Balaji Krishnan, Muralidhar L. Hegde, Michael Weinfeld, Gourisankar Ghosh, Tapas Hazra

**Author notes:** Contributed equally (Anirban Chakraborty, Santi M. Mandal). Address correspondence to: Tapas Hazra, Department of Internal Medicine-Pulmonary, Critical Care & Sleep Medicine, University of Texas Medical Branch, Galveston, TX 77555, USA; Gourisankar Ghosh, Department of Chemistry and Biochemistry, University of California San Diego, LA Jolla, California 92093, USA.

## Abstract

Several reports have indicated that impaired mitochondrial function contributes to the development and progression of Huntington’s disease (HD). Mitochondrial genome damage, particularly DNA strand breaks, is a potential cause for its compromised functionality. Here we show that the activity of polynucleotide kinase 3’-phosphatase (PNKP), a critical DNA end-processing enzyme, is significantly decreased in the mitochondrial extract of HD patients’ brains due to a lower level of fructose-2,6 bisphosphate (F2,6BP), a biosynthetic product of 6-phosphofructo-2-kinase fructose-2,6-bisphosphatase 3 (PFKFB3). Such decrease in PNKP activity leads to persistent DNA strand breaks that are refractory to subsequent steps for repair completion. Both PFKFB3 and F2,6BP, an allosteric modulator of glycolysis, are also present in the mitochondria and PFKFB3 is part of a mitochondrial DNA repair complex containing HTT, PNKP, DNA Pol γ (POLG) and Lig IIIα. Notably, PNKP binds F2,6BP (Kd= 525±25 nM) and utilizes it as a cofactor. The levels of both F2,6BP and PFKFB3 are significantly decreased in the mitochondrial extract of HD mouse striatal neuronal cells and patients’ brain. Activity of PNKP is thus severely decreased in the mitochondrial extract; however, addition of F2,6BP restored its activity. Moreover, supplementation of F2,6BP in HD cells restored PFKFB3 level, mitochondrial genome integrity and partially restored mitochondrial membrane potential, mitochondrial respiration and prevented pathogenic aggregate formation. We also observed that supplementation with F2,6BP restored mitochondrial genome integrity in an HD *Drosophila* model. Our findings, therefore, suggest that F2,6BP-mediated restoration of PNKP activity could have a profound impact in ameliorating neurodegenerative symptoms in HD.

**Significance:** We reported earlier the loss of PNKP activity in the nuclear extracts from HD patients’ brain. However, a glycolytic metabolite, F2,6BP, can restore PNKP activity and rescue organismal phenotypes in HD fly models. As PNKP is present in mitochondria and several reports indicate that mitochondrial dysfunction contributes to HD, we therefore analyzed PNKP activity in the mitochondrial extract. Surprisingly, we found that PFKFB3 and its product, F2,6BP are present in mitochondria, but significantly low in patients’ brains. Exogenous addition of F2,6BP restored PNKP activity in patients’ brain mitochondrial extract. Moreover, supplementing F2,6BP in HD cells and fruit flies restored mitochondrial genome integrity suggesting maintaining adequate intracellular F2,6BP levels is critical for proper functionality of PNKP and thereby of brain health.

## Introduction

Huntington’s disease (HD), a devastating hereditary neurological disorder, is caused by toxic expansion of the polyglutamine (polyQ) stretch in the N-terminus of huntingtin protein (HTT)^1^. Although, HD is an autosomal, monogenic disease, the underlying mechanisms that contribute to the death of brain cells in HD are still elusive. Genome-wide association studies showed a strong correlation between DNA repair deficiency and the age-dependent onset of HD^2,3^. In addition to nuclear genome damage biology, recent advances in mitochondrial research have implicated previously unanticipated roles of this organelle in human diseases, particularly aging and neurological disorders, including HD^4-7^. Mitochondrial DNA (mtDNA) damage is a potential cause of mitochondrial dysfunction and can have a significant impact on cellular health in HD^8-9^. The mitochondrial genome is subjected to continuous insult by endogenous reactive oxygen species (ROS) because of its proximity to the site of ROS generation via mitochondrial electron transport system complexes. Furthermore, mtDNA lacks protective histones, unlike nuclear DNA (nDNA), and so it is more susceptible to such oxidative damage^10^. Moreover, mitochondria have a high rate of transcription and importantly, most of the mitochondrial genome is transcribed. Thus, mitochondrial DNA encoded core functions would be compromised due to deficient mitochondrial genome repair in HD^9,11-13^.

To protect mtDNA from ROS induced oxidative damage, a range of DNA repair mechanisms have evolved including base excision repair (BER) and single strand break repair (SSBR)^13^. Polynucleotide kinase 3’-phosphatese (PNKP) is a major enzyme for processing both “non-ligatable” 3’-phosphate (3’-P) and 5’-OH termini at strand breaks in mammalian genomes and thus, it is involved in multiple repair pathways including BER and SSBR^14-15^. Several recent reports, including ours, have documented an association of PNKP deficiency with neurological/developmental disorders^16-20^. However, the underlying biochemical bases of such phenotypes, which are exclusive to the nervous system, are not well understood. We have recently reported that wild-type HTT plays an important role in DNA repair most likely by providing a platform for the assembly of a novel transcription-coupled DNA repair (TCR) complex in nuclei that includes, RNA polymerase II (RNAPII), PNKP and other DNA repair proteins^21-22^. This specialized protein complex repairs DNA lesions during transcription to maintain genome integrity of the neurons, preserve their function and by doing so, extend their survival. We have further demonstrated cellular toxicity due to the loss of DNA repair (3’-phosphatase) activity of PNKP, but not its protein level, in HD brain extracts, leading to accumulation of DNA SBs, including double strand breaks^22^. Most importantly, we have observed that PNKP interacts with the nuclear isoform of a glycolytic enzyme, 6-phosphofructo-2-kinase fructose-2,6-bisphosphatase 3 (PFKFB3). Depletion of PFKFB3 markedly reduced PNKP activity without changing its protein level. Notably, the levels of both PFKFB3 and its product fructose-2,6 bisphosphate (F2,6BP), an allosteric modulator of glycolysis, are significantly lower in the nuclear extracts of post-mortem HD patients’ brains. Supplementation of F2,6BP in HD mouse striatal neuronal cells and in HD fruit flies fully restored nuclear genome integrity and functionality^22^, suggesting F2,6BP to be a positive regulator of PNKP activity *in vivo*.

The repair of ROS-induced DNA damage in the mitochondrial genome is well characterized. We and others have shown the presence of DNA glycosylases, NEIL1/NEIL2 and PNKP in the mitochondria^23-25^. NEIL1 and NEIL2 initiate BER by their combined DNA glycosylase and AP lyase activities and generate SBs with 3’-P termini, a substrate for PNKP. In addition to impeding DNA repair, 3’-P termini can stall elongating RNA polymerases, leading to DNA damage response via p53 activation^26^. Thus, processing of such “non-ligatable” 3’-P-containing DNA termini is essential for repair progression and efficient transcription even within mitochondria. Since PNKP, the major 3’-phosphatase in mammalian cells^27-28^, is also present in mitochondria^24-25^, we, therefore, asked whether the activity of PNKP is compromised in mitochondria of HD patients and whether PFKFB3 or F2,6BP has any role in mtDNA repair, similar to what we reported earlier for nuclear DNA repair^22^. Surprisingly, we observed that both PFKFB3 and F2,6BP localize to mitochondria and their levels are significantly low in HD patients’ mitochondrial extract, leading to abrogated mtPNKP activity. Supplementing F2,6BP in HD-mouse striatal neuronal cells and a *Drosophila* HD model system^29^ restored mitochondrial genome integrity. Altogether, our data provides a strong rationale for exploring the therapeutic potential of F2,6BP or its analog in HD and related pathologies.

## Results and discussion

### Mitochondrial PNKP activity is decreased in HD

Our previous studies have demonstrated that nuclear PNKP activity is diminished in HD mouse models^21^ and post-mortem brain tissues from HD patients^22^. Given that PNKP is known to translocate to mitochondria^24^, we investigated whether its activity is also compromised in mitochondria by assessing the 3’-phosphatase activity of PNKP in the mitochondrial extracts prepared from the frontal cortex of post-mortem HD brains vs healthy controls. We indeed observed a marked reduction of PNKP activity in HD samples **(Fig. 1A, lanes 6-9 vs lanes 2-5)**. Mitochondria are important sources of cellular reactive oxygen species (ROS), essential byproducts generated from leaked electrons and molecular oxygen. While ROS are natural signaling molecules regulating various biological processes, excessive mitochondrial ROS (mtROS) induce oxidative stress, damaging essential biomolecules, including mitochondrial DNA (mtDNA). Therefore, we also assessed the mtROS level in diseased and healthy control tissue. We indeed observed a significantly higher level of ROS in HD patients **(Fig. 1B)**.

**Figure 1:**
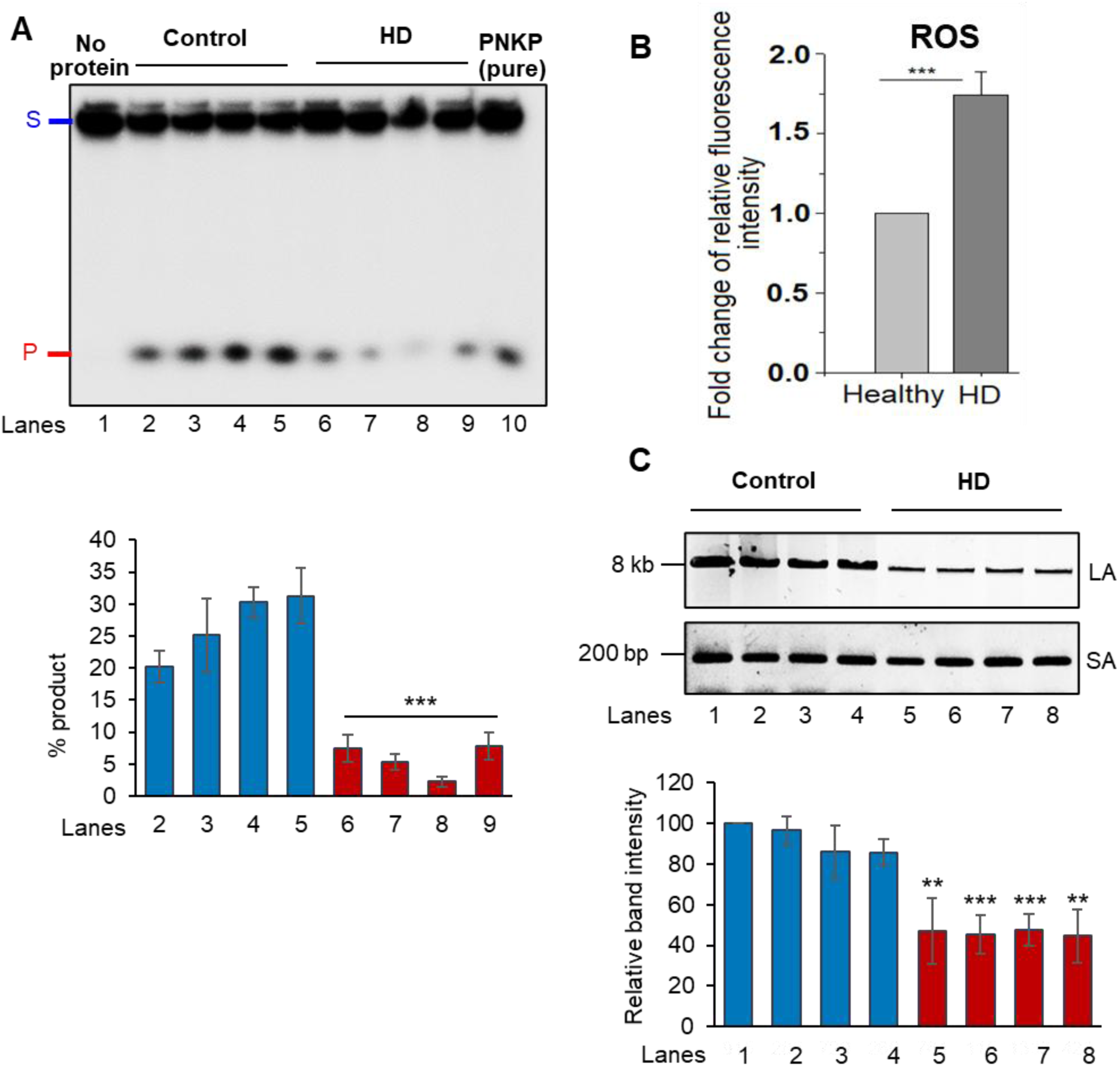
**A. Upper panel:** 3’-phosphatase activity of PNKP in the mitochondrial extract (250 ng) of frontal cortex from healthy normal control (lanes 2-5) vs. age-matched HD patients (lanes 6-9) measured by the release of free phosphate from a radiolabeled 3’-phosphorylated substrate. Lane 1: substrate only. Lane 10: Purified PNKP (2 ng). S: Substrate and P: Released phosphate. **Lower panel:** Quantitation of the % released phosphate in the indicated lanes (n=3); error bars represent mean ± SD. ***P<0.005 between Control vs. HD patient. **B.** Bar diagram represents mtROS level as relative fluorescence intensity with healthy control arbitrarily chosen as unity (n=14; ***P<0.005). **C. Upper panel:** Representative agarose gel image of long (LA) and short (SA) amplicon of the mitochondrial fragment from genomic DNA of postmortem frontal cortex of age-matched healthy normal (control) (lanes 1-4) and HD patients (lanes 5-8). **Lower panel:** The normalized relative band intensities are represented in the bar diagram with one control sample arbitrarily set as 100. The damage accumulation in HD patients significantly increased (***P < 0.005; **P < 0.01).

Since PNKP is an essential DNA end-processing enzyme, we next examined accumulation of DNA SBs in the mitochondrial genome by a long amplicon (LA)-qPCR-based assay using mitochondrial genome specific primers. In this assay, a relative decrease in the PCR product of the long amplicon (∼8-10 kb) vs the short amplicon (∼250 bp) suggests increased DNA damage, as a higher number of lesions in the longer templates impedes PCR amplification. We observed significant accumulation of DNA SBs in the mitochondrial genome of HD brains **(Fig. 1C, lanes 5-8 vs lanes 1-4)** concurrent with higher level of mtROS and compromised mtPNKP activity. In summary, our results indicate that mtDNA also accumulates damage due to a significant reduction in mtPNKP activity in HD pathology, consistent with our earlier report showing significant increase in nuclear genome damage in HD patients^22^.

### Diminished PFKFB3 and F2,6BP levels in HD patients disrupt mitochondrial PNKP activity

Given that PNKP levels are similar between HD patients and healthy control samples, and that F2,6BP acts as a cofactor of PNKP to maintain its activity in the nucleus, we postulated that a similar regulatory mechanism is present in mitochondria. To assess the role of PFKFB3 and its product F2,6BP in mitochondrial DNA repair, we first tested whether PFKFB3 is localized in mitochondria in HEK293 cells. Indeed, we detected significant amounts of PFKFB3 in the purified mitochondrial extract of HEK293 cells **(Fig. 2A)**. We further confirmed the presence PFKFB3 in the mitochondrial extract from WT mouse striatal neuronal (Q-7) cells and post-mortem frontal cortex from healthy controls **(Fig. 2B, lane 1; Fig. 2C, lanes 1-4)**. Since PFKFB3 is present within mitochondria, we tested for its association with mtPNKP and other repair proteins by immunopulldown of PFKFB3 from mitochondrial extract of Q-7 cells using an anti-PFKFB3 antibody. We observed PNKP, mitochondrial DNA polymerase γ (POLG) and HTT in the PFKFB3 immunocomplex **(Fig. 2D)**, confirming PFKFB3 as a part of a mitochondrial DNA repair complex. Absence of RNAPII in the mitochondrial extract and immunocomplex provided evidence in favor of mitochondria specific complex formation. To further validate these findings, we performed indirect immunofluorescence and co-staining of mitochondria and nuclei (by Mito tracker green and DAPI, respectively) which confirmed localization of PFKFB3 in both the mitochondria and nucleus of Q-7 cells **(Fig. 2E)**. We further observed that mitochondrial integrity was severely compromised in striatal neuronal cells from HD mouse with expanded polyQ repeats (Q-111), with a significant reduction in PFKFB3 levels **(Fig. 2F, 2H)**. Western blot analyses also revealed a significant decrease in PFKFB3 levels in the mitochondrial extracts of Q-111 cells **(Fig. 2B, lane 2 vs lane 1)** and HD post-mortem patients’ brains **(Fig. 2C, lanes 5-8 vs lanes 1-4)**. However, the level of PNKP was similar in both Q-111 cells and HD patient extracts compared to respective controls **(Fig. 2B and 2C)**. consistent with our previous findings^21,22^. These observations suggest that the DNA strand break repair components are active in mitochondria of healthy cells/tissues but are compromised under pathogenic conditions. The presence of PFKFB3 in the nucleus and mitochondria further emphasizes its additional function in the DNA repair pathway beyond its known role in glycolysis.

**Figure 2:**
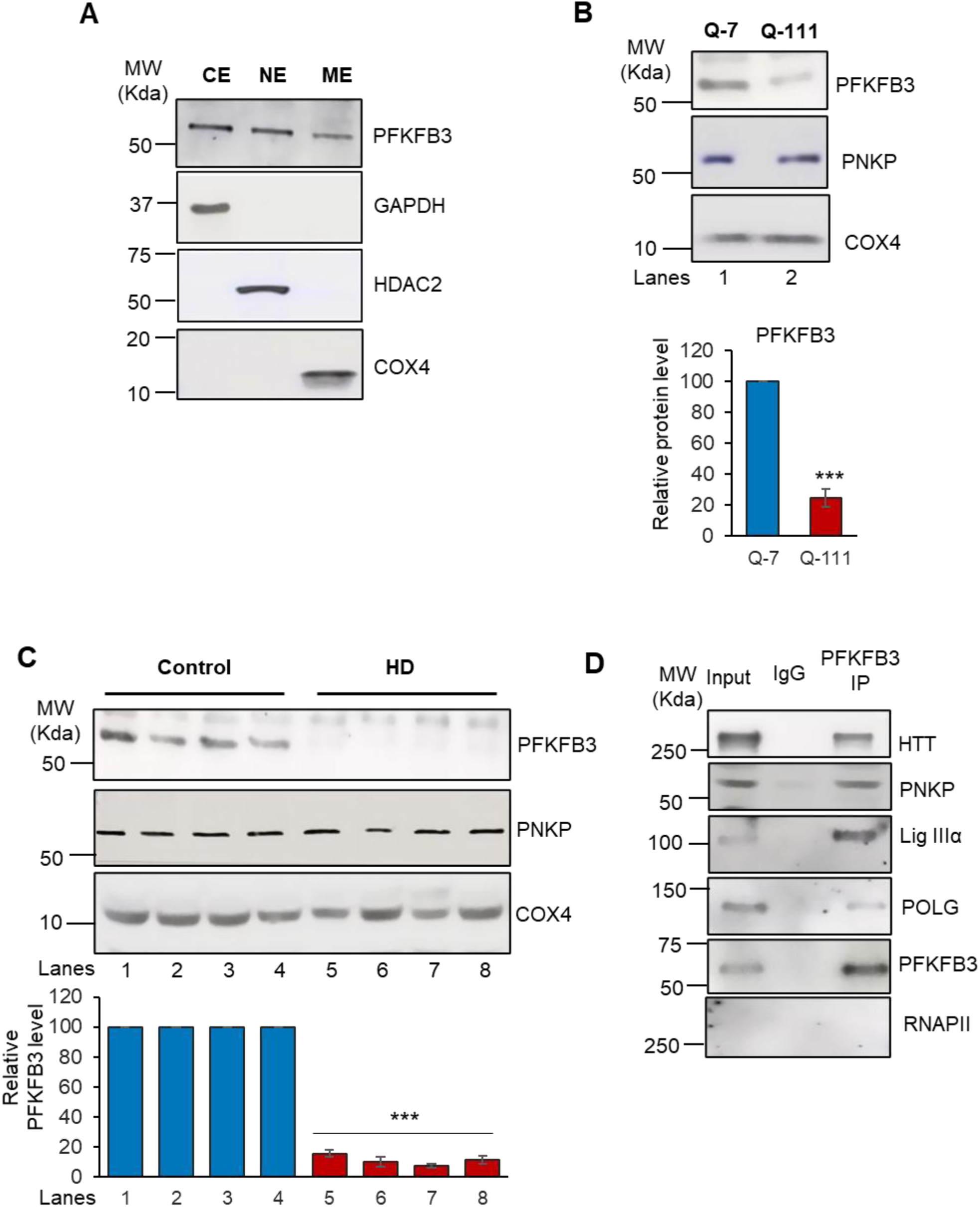
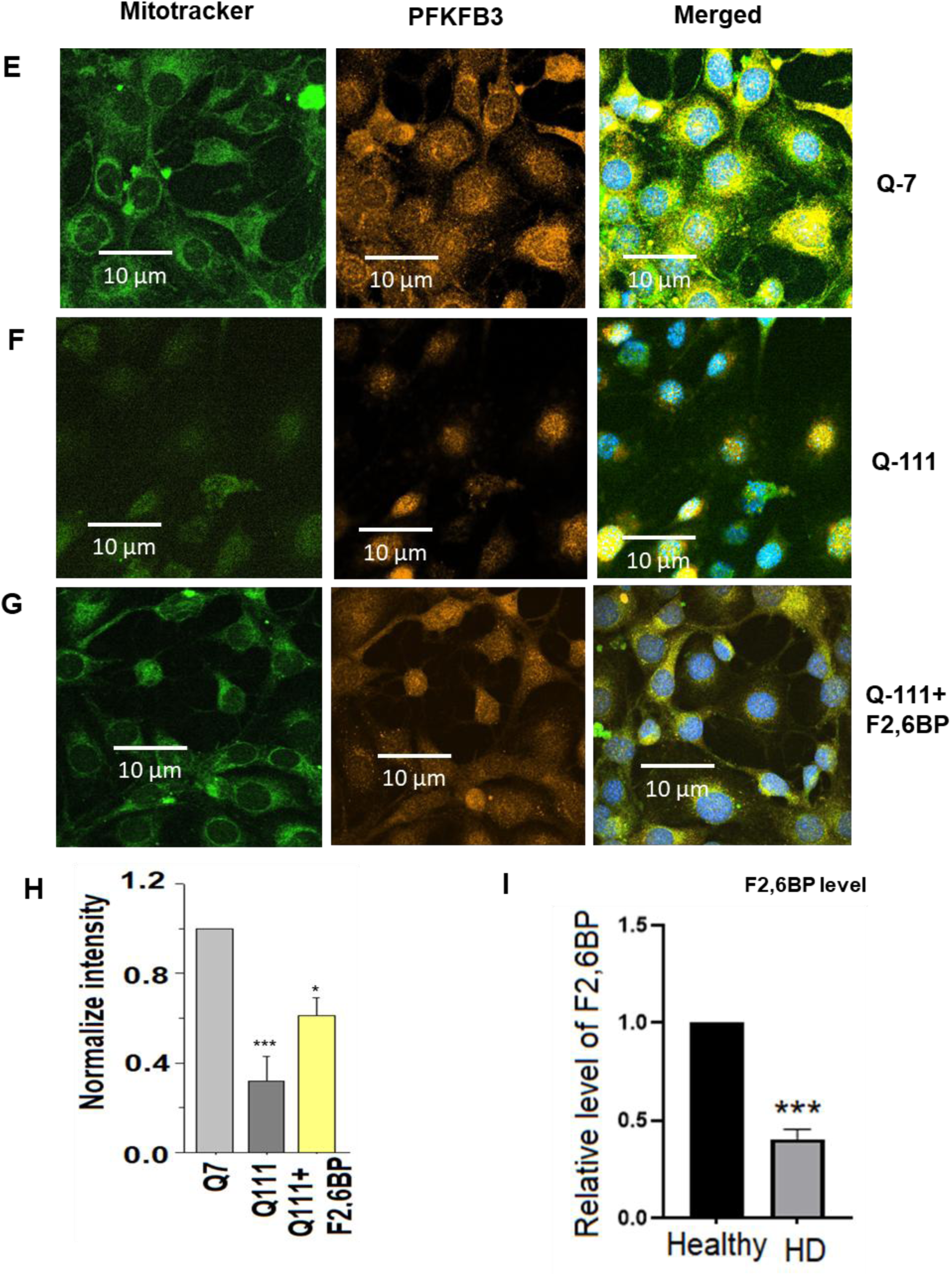
**A.** Western blot showing the relative levels of PFKFB3 in the cytosolic (CE), nuclear (NE) and mitochondrial (ME) extract of HEK293 cells. GAPDH: cytosolic loading control; HDAC2: nuclear loading control. COX4: mitochondrial loading control. **B.** Western blot showing the relative levels of PNKP and PFKFB3 in the mitochondrial extract of Q-7 and Q-111 cells. COX4: mitochondrial loading control. **Lower panel:** Quantitation of the relative PFKFB3 levels after normalization with loading control COX4 (n=3, ***P<0.005). **C.** Western blot showing the relative levels of PNKP and PFKFB3 in the mitochondrial extract of HD patients vs. age-matched control subjects’ frontal cortex. COX4: mitochondrial loading control. **Lower panel**: Quantitation of the relative PFKFB3 levels after normalization with loading control COX4 (n=3, ***P<0.005). **D.** Benzonase-treated mitochondrial extracts from Q-7 cells were immunoprecipitated (IP’d) with anti-PFKFB3 antibody (PFKFB3 IP); or control IgG and tested for the presence of associated proteins (shown in the right) using specific Abs. **E-G.** Q-7 **(E)** and Q-111 **(F)** cells were first stained with MitoTracker dye, which marks mitochondria in green **(left panels)**, followed by fixation and staining for PFKFB3 using a secondary antibody conjugated to Alexa Fluor 568 **(middle panels)**. The overlap of these signals appears yellow in merged images **(right panels)**, confirming PFKFB3’s mitochondrial localization. **(G)** Represents Q-111 cells after treatment with F2,6BP (50 µM), showing improved co-localization in the merged image suggesting that F2,6BP may help restore mitochondrial integrity and PFKFB3 levels in Q-111 cells. Nuclei are counterstained with DAPI. **H.** Quantitation of the PFKFB3 level (***P<0.005; *P<0.05). **I.** Bar diagram showing the relative levels of F2,6BP in the mitochondrial extract of control vs. HD patients’ frontal cortex (n=14, ***P<0.005).

We next examined if F2,6BP is present in the mitochondria of healthy individuals and if its level is reduced in the post-mortem HD patients in line with low PFKFB3 levels and PNKP activity. Not only did we detect F2,6BP in the mitochondrial extract of healthy subjects, but we also found that the level of F2,6BP was nearly 2.5-fold lower in the mitochondrial extracts of patient samples (**Fig. 2I**; Left Panel). Since the level of mtPNKP protein remained comparable in HD patients and their age-matched controls, consistent with our previous observations in nuclei, the reduction in mtPNKP activity is therefore due to lower levels of F2,6BP and not due to decrease in the level of PNKP^22^. These results convincingly suggest that an endogenous metabolite F2,6BP, a well-known potent activator of glycolysis, also plays an important additional role in maintaining activity of an essential DNA repair enzyme and thereby contributes to cellular health.

### Dysregulation of mitochondrial function in HD

To gain further insight into mitochondrial quality and functionality, we measured the mitochondrial membrane potential, a key indicator of mitochondrial health. For this purpose, we utilized tetramethyl rhodamine methyl ester (TMRM) that is readily sequestered in healthy cells with functional mitochondria, emitting a red-orange, fluorescent signal. Flow cytometry (FACS) analysis revealed a significant decrease in membrane potential in the Q-111 cells relative to Q-7 cells **(Fig. 3A-C)**.

**Figure 3:**
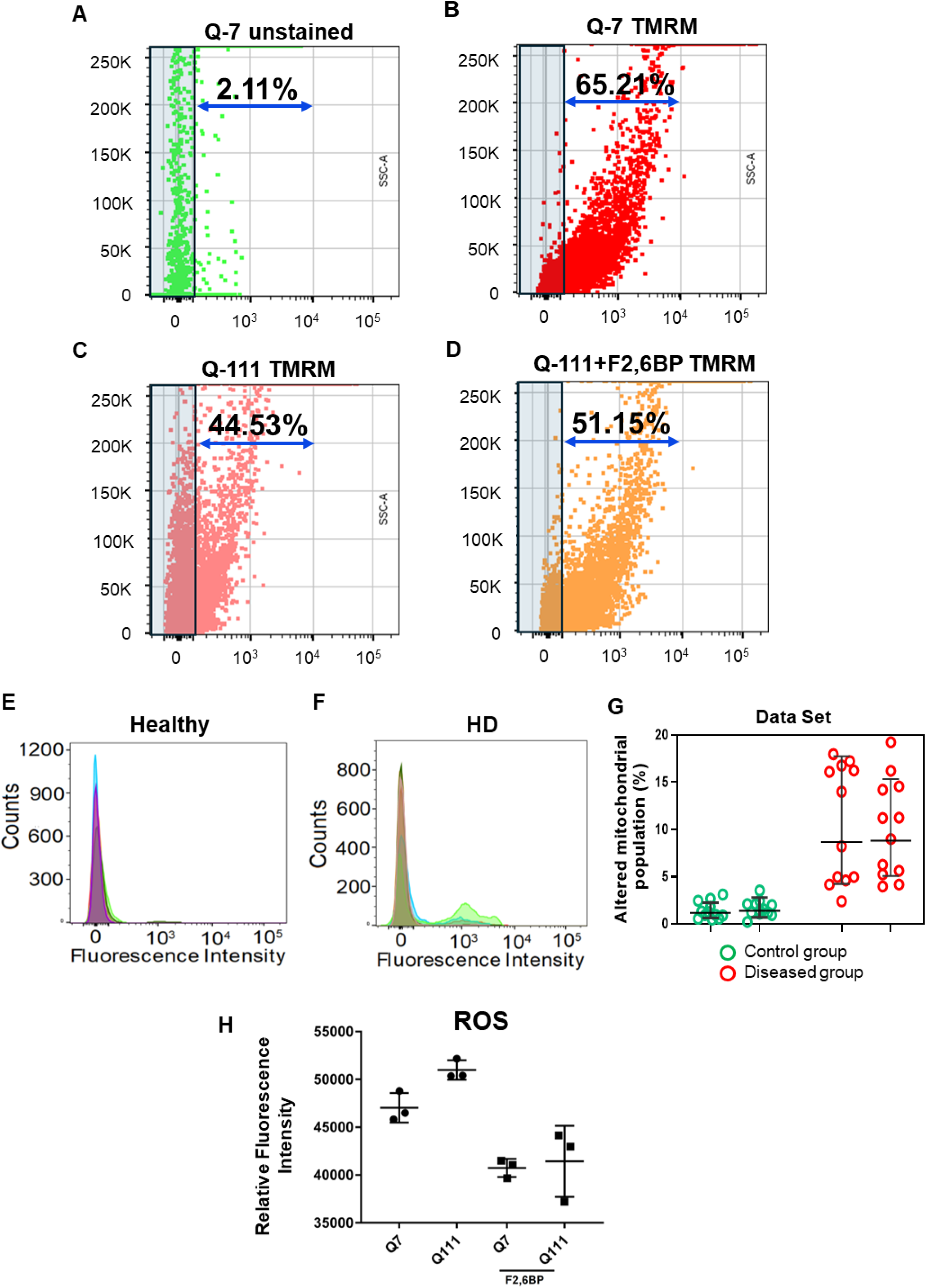
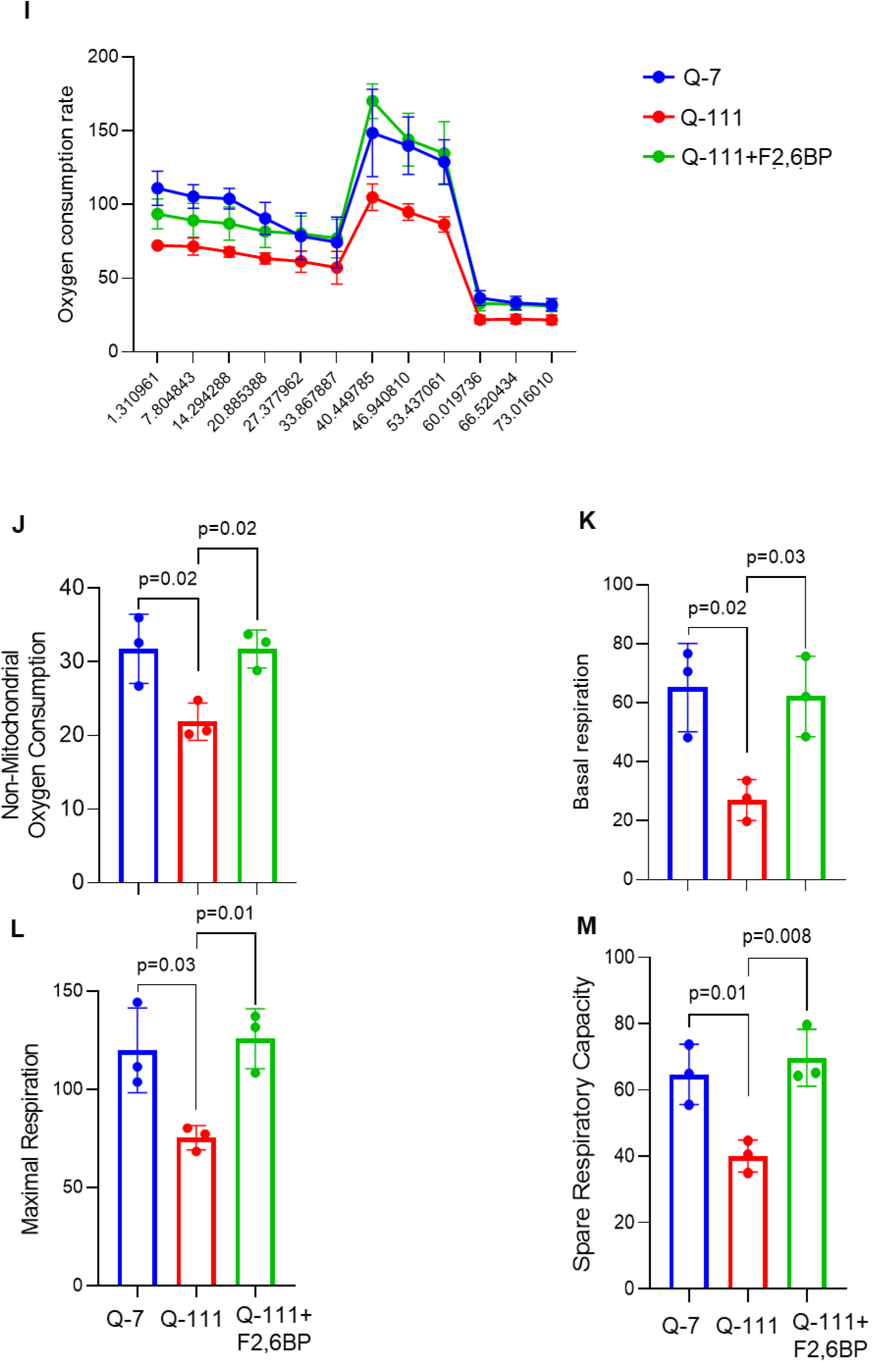
**A-D:** Flow cytometry scatter plot analysis illustrates changes in mitochondrial membrane potential in Q-7 cells without TMRM **(A)**, Q-7 cells with TMRM **(B)**, Q-111 cells with TMRM **(C)**, and F2,6BP-treated (100 µM) Q-111 cells with TMRM **(D)**. Fluorescence intensity of TMRM was measured using the PE channel. **(E-G)** Isolated mitochondria from both healthy and HD tissues were analyzed by FACS using the MitoView™650 dye on the PE channel, revealing distinct populations with different mitochondrial percentages. Mitochondria from diseased tissues **(F)** demonstrated a separate population compared to mitochondria from healthy control tissues **(E)**. Quantitative results from duplicate analyses are summarized in the bar graph **(G)** and are expressed as mean ± SE. **H.** MtROS level in Q-7 and Q-111 cells ± F2,6BP (100 µM) in terms of relative fluorescence intensity. **I-M:** Mitochondrial respiration comparison between Q-7, Q-111 and Q-111 cells supplemented with 200 µM F2,6BP by seahorse assay. Oxygen consumption rate (OCR) **(I)** was determined throughout the mitochondrial respiration test. Non-mitochondrial oxygen consumption **(J)** was determined by measuring the difference between total oxygen consumption and antimycin A and rotenone treatment-induced reduction in oxygen consumption, maximal respiration **(L)** was expressed as difference between oxygen consumption following mitochondria uncoupling by FCCP and rotenone, antimycin A treatment and spare respiratory capacity **(M)** was determined by subtracting basal respiration **(K)** from maximal respiration. The data represent mean ± s.e.m. from three independent experiments.

Additionally, we used MitoView™650, a mitochondria specific dye which enables real-time visualization of mitochondrial morphology, distribution, and dynamics in live cells without relying on membrane potential. This dye has an excitation/emission spectrum of 644/670 nm, compatible with the Cy®5 channel. Pure mitochondria were isolated from post-mortem HD brain frontal cortex vs healthy controls, stained with MitoView™650, and analyzed using the 670 nm channel. A single uniform population of mitochondria was found in healthy controls; however, heterogeneous mitochondrial populations were observed in HD samples **(Fig. 3E-G)**, which may be attributed to some irregularities in mitochondrial function and health associated with HTT aggregation or reduced level of F2,6BP. Hence, our data clearly indicates that decreased levels of mitochondrial PFKFB3/F2,6BP in HD are associated with compromised mitochondrial health and function. Consistent with our results in the HD patients, we observed significantly higher level of mtROS in Q-111 cells **(Fig. 3H)**, indicating pathological consequences.

To assess mitochondrial function further, we measured cellular respiration in Q-7 vs Q-111 cells **(Fig. 3I-M)**. These measurements were carried out using the Seahorse XF Cell Mito Stress Test protocol, both under baseline conditions and after the application of mitochondrial inhibitors. Our observations indicated a significant reduction in oxygen consumption rate (OCR) in Q-111 cells **(Fig. 3I)**. We also observed significantly less non-mitochondrial oxygen consumption **(Fig. 3J)**, and in their basal, maximal, and spare respiration capacities **(Fig. K-M)**. Taken together, these results suggest that a pool of endogenous PFKFB3 is localized in mitochondria and that the level of PFKFB3 and its metabolic product, F2,6BP is less in Q-111 HD cells that correlates with compromised mitochondrial function, such as mitochondrial membrane potential and respiration.

### Exogenous F2,6BP restores mitochondrial PNKP activity by direct binding

Given that F2,6BP enhanced PNKP activity in the HD patient brain-derived nuclear extracts^22^, we investigated whether F2,6BP could similarly restore PNKP activity in mitochondrial extracts. We synthesized F2,6BP *in vitro* biochemically using recombinant PFKFB3 followed by purification using ion exchange chromatography. Electrospray ionization (ESI) mass-spectrometry showed the purity of F2,6BP **(Fig. 4A)**. We then determined the binding affinity of PNKP for F2,6BP by monitoring changes in intrinsic tryptophan fluorescence of PNKP upon binding to F2,6BP. Addition of F2,6BP (400 nM) to PNKP resulted in partial quenching of the tryptophan fluorescence of PNKP at 340 nm, when excited at 295 nm with no change in the emission maximum **(Fig. 4B)**. Binding affinity (Kd= 525 ± 25 nM) was determined by following fluorescence quenching (a measure of ligand binding) as a function of F2,6BP concentration **(Fig. 4C)**. A representative plot of relative fluorescence intensities versus concentration of F2,6BP is shown in **Fig. 4C (inset)**.

**Figure 4:**
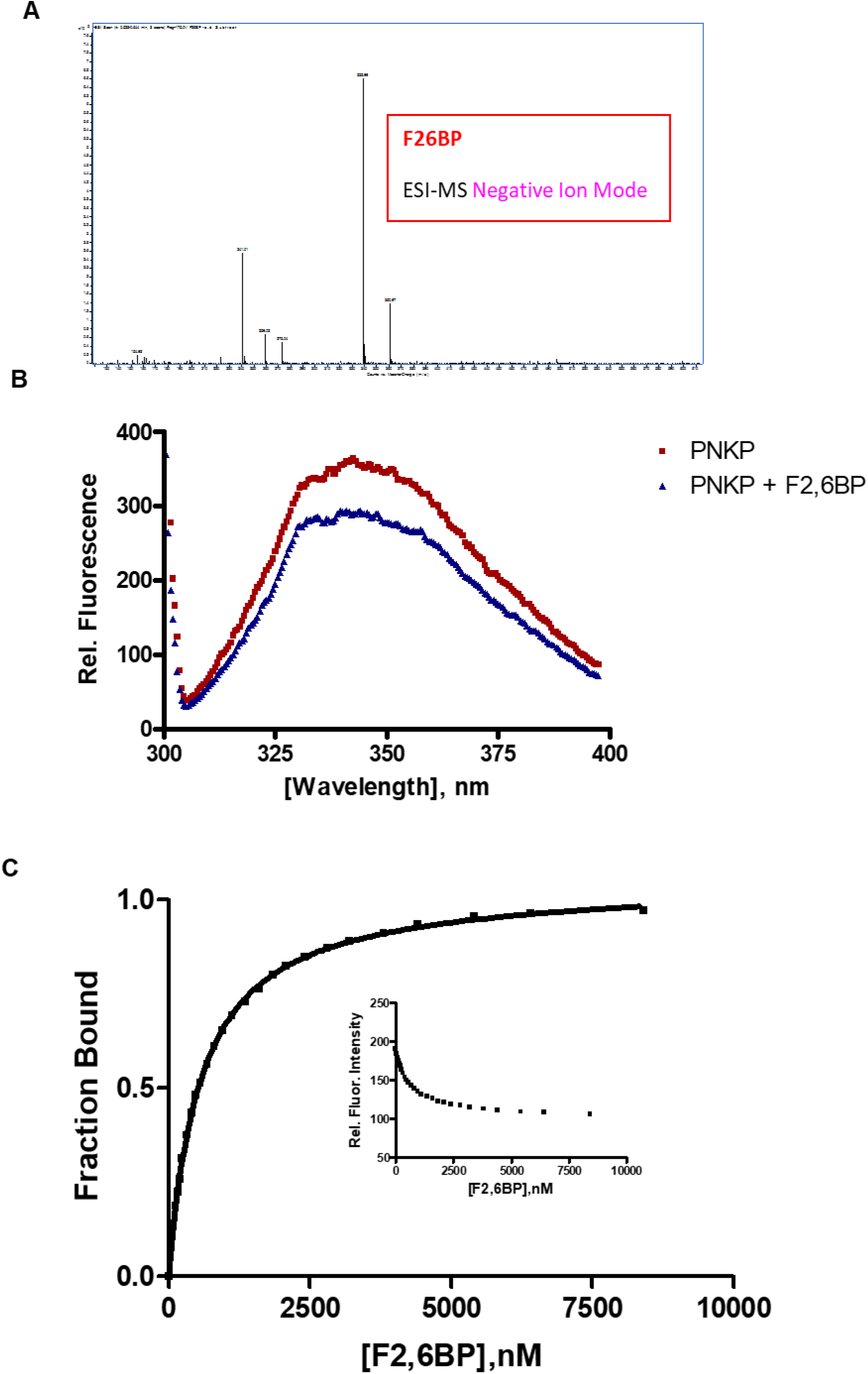
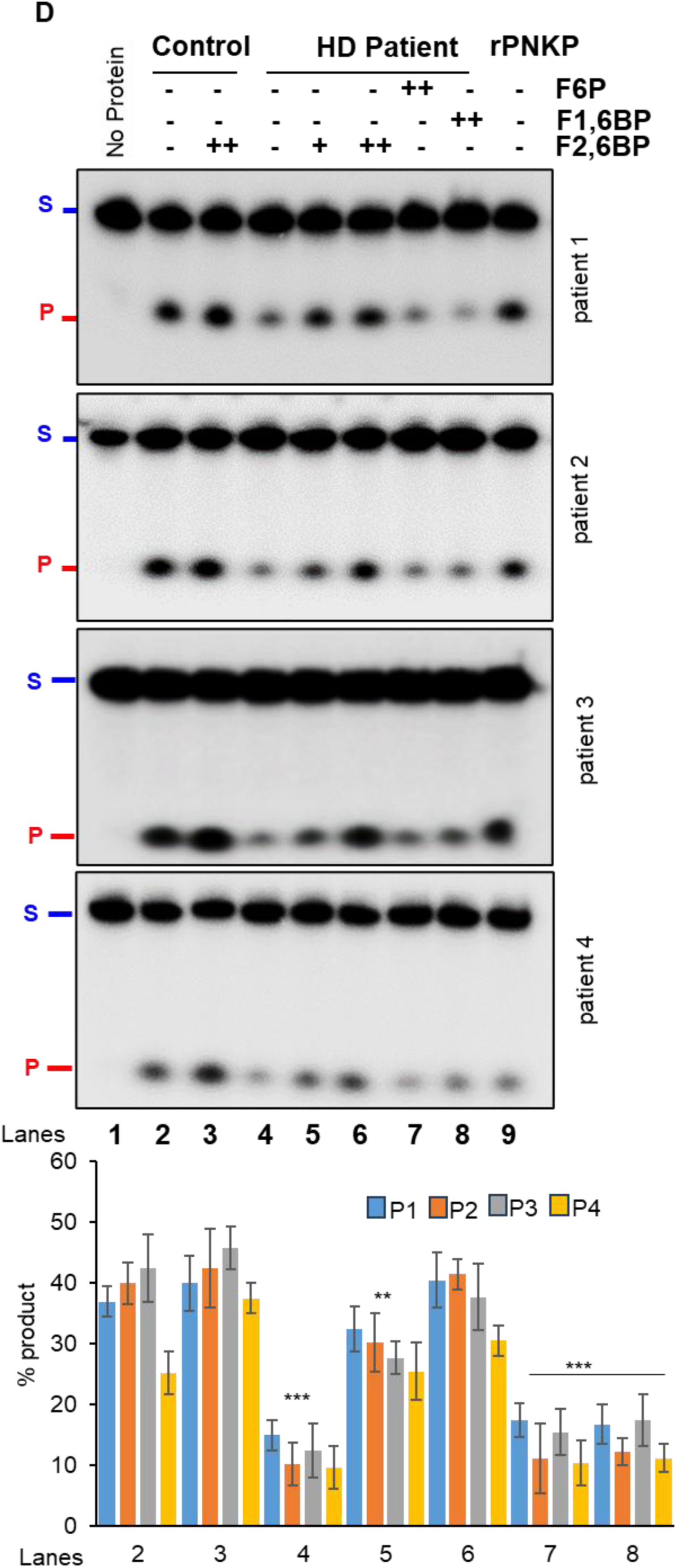
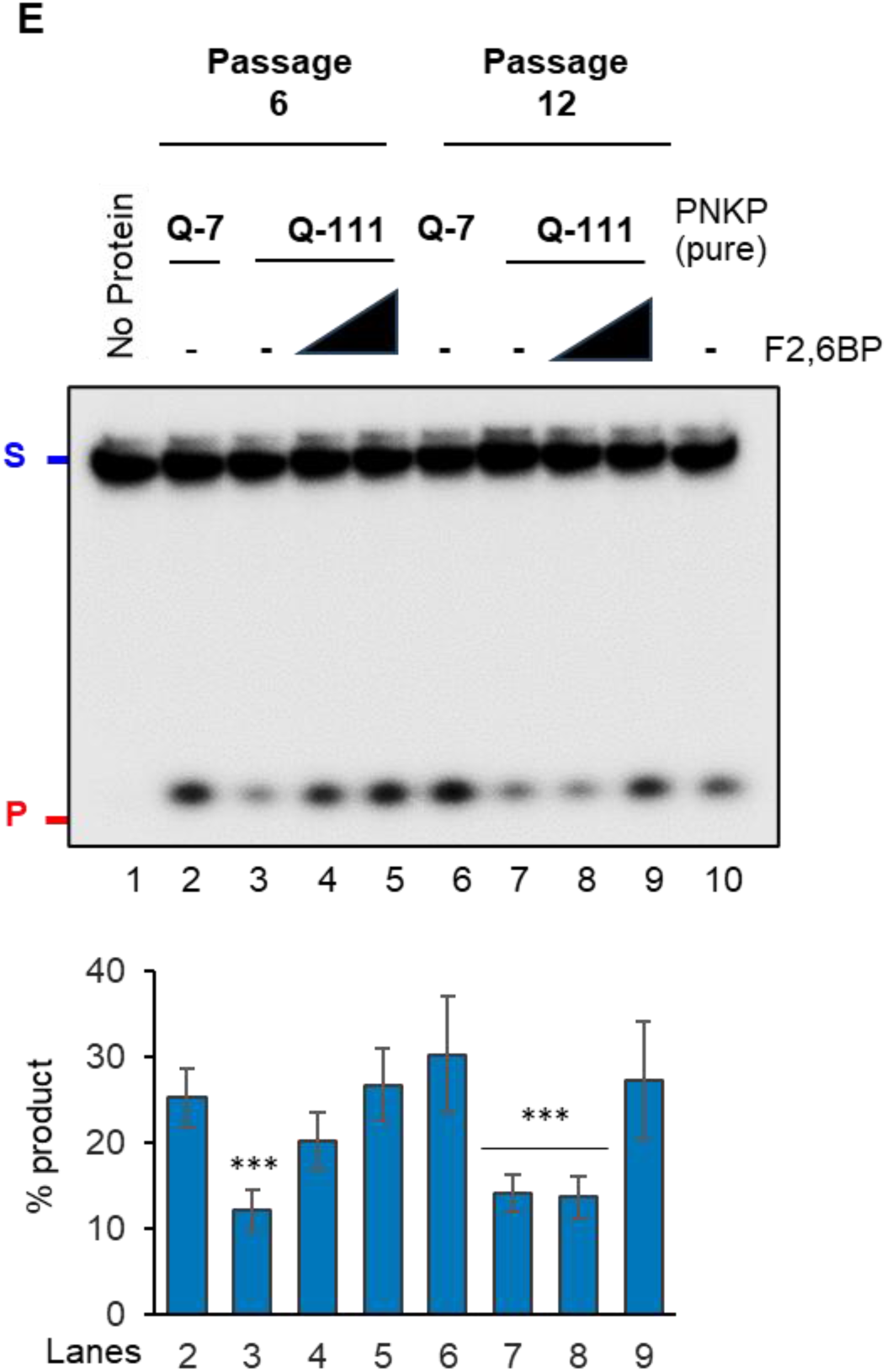
**A.** ESI-MS analysis of the purified F2,6BP in negative ion mode showing a prominent peak at m/z 338.99, which corresponds to the deprotonated molecular ion (M-H]- of F2,6BP. **B.** Fluorescence emission spectra of PNKP excited at 295 nm in the absence and the presence of F2,6BP. **C.** Fluorescence titration of PNKP versus F2,6BP. The protein (50 nM) was excited at 295 nm, and the fluorescence intensity was monitored at 340 nm (see inset) at room temperature. The fraction bound (i.e. relative fluorescence quenching) versus F2,6BP concentration is plotted. **D. Upper panel:** Representative gel images (four different patients vs healthy control) showing the 3’-phosphatase activity of PNKP in the post-mortem brain (frontal cortex) mitochondrial extract of healthy normal control (lane 2) or supplemented with F2,6BP (lane 3, 50 µM), and age/gender-matched HD patient (lane 4) or supplemented with F2,6BP (lanes 5-6, 25 and 50 µM) or F6P (lane 7, 50 µM) or F1,6BP (lane 8, 50 µM). Lane 1: substrate only. Lane 9: purified PNKP (1-2 ng). S: Substrate and P: Released phosphate. **Lower panel:** Quantitation of the % released phosphate in the indicated lanes (n=3, ***P<0.005; **P<0.01). **E. Upper panel:** Representative gel image showing the 3’-phosphatase activity of PNKP in the mitochondrial extract of Q-7 cells (lanes 2, 6) and Q-111 cells (lanes 3, 7) or Q-111 cells supplemented with F2,6BP (lanes 4-5 and 8-9, 25-50 µM). Lane 1: substrate only. Lane 10: purified PNKP (1-2 ng). S: Substrate and P: Released phosphate. Cells from two different passages are used (P6 and P12). **Lower panel:** Quantitation of the % released phosphate in the indicated lanes (n=3, ***P<0.005, between Q-7 and Q-111 cells).

We next investigated the effect of F2,6BP in restoring activity of PNKP in the mitochondrial extracts. As expected, exogenous F2,6BP restored the 3’-phosphatase activity of PNKP in a dose-dependent manner in mitochondrial extracts isolated from post-mortem HD brains (frontal cortex) across different age and gender groups **(Fig. 4D, lanes 5-6 vs lane 4)**. However, related natural metabolites, neither fructose-6-phosphate (F6P) nor fructose 1,6-bisphosphate (F1,6BP) could restore PNKP activity **(Fig. 4D, lanes 7 and 8)**, highlighting the specificity of F2,6BP in reactivating PNKP. In our previous study, we found compromised PNKP activity in the nuclear extract of Q-111 cells compared to WT cells (Q-7)^22^. Therefore, we assessed PNKP activity in the mitochondrial extracts of Q-7 vs Q-111 cells and observed a significant reduction in 3’-phosphatase activity in Q-111 cells **(Fig. 4E, lane 3 vs lane 2; lane 7 vs lane 6)**. A similar dose-dependent restoration by F2,6BP was observed in mitochondrial extracts from Q-111 cells **(Fig. 4E, lanes 4-5 vs lane 3; lanes 8-9 vs lane 7)**, underscoring the essential role of F2,6BP as a co-factor, in enhancing PNKP-mediated DNA repair in mitochondria.

To further evaluate the potential of exogenous F2,6BP in restoring DNA repair deficiency at the cellular level, we assessed *in-cell* rescue of genome integrity and functionality in Q-111 cells compared to WT Q-7 cells^30^. F2,6BP was delivered following an optimized protocol^22^ and allowed 72 or 96 hours to alleviate mtDNA damage-induced cellular toxicity. As a control, Q-111 cells were either mock transfected or transfected with F1,6BP. The restoration of 3’-phosphatase activity of PNKP was observed in a time-dependent manner exclusively in the mitochondrial extracts of F2,6BP-transfected cells **(Fig. 5A, lanes 6-7 vs lane 4)**, but not in cells transfected with either F1,6BP or the mock-transfected cells **(Fig. 5A, lanes 8-9 or lane 5)**. We also observed significant restoration of mitochondrial genome integrity following treatment with F2,6BP, concurrent with rescue of mtPNKP activity **(Fig 5B, lanes 5-6 vs lane 3)**. In our earlier study, we observed a significant increase in the Q-111 cell count following F2,6BP treatment, comparable to control Q-7 cells, contrary to the decreased count under mock treatment condition^22^. Therefore, our study further indicates that apart from nuclear DNA repair, repair of mtDNA substantially contributes to the rescue of the diseased cells from neurotoxicity and apoptosis.

**Figure 5:**
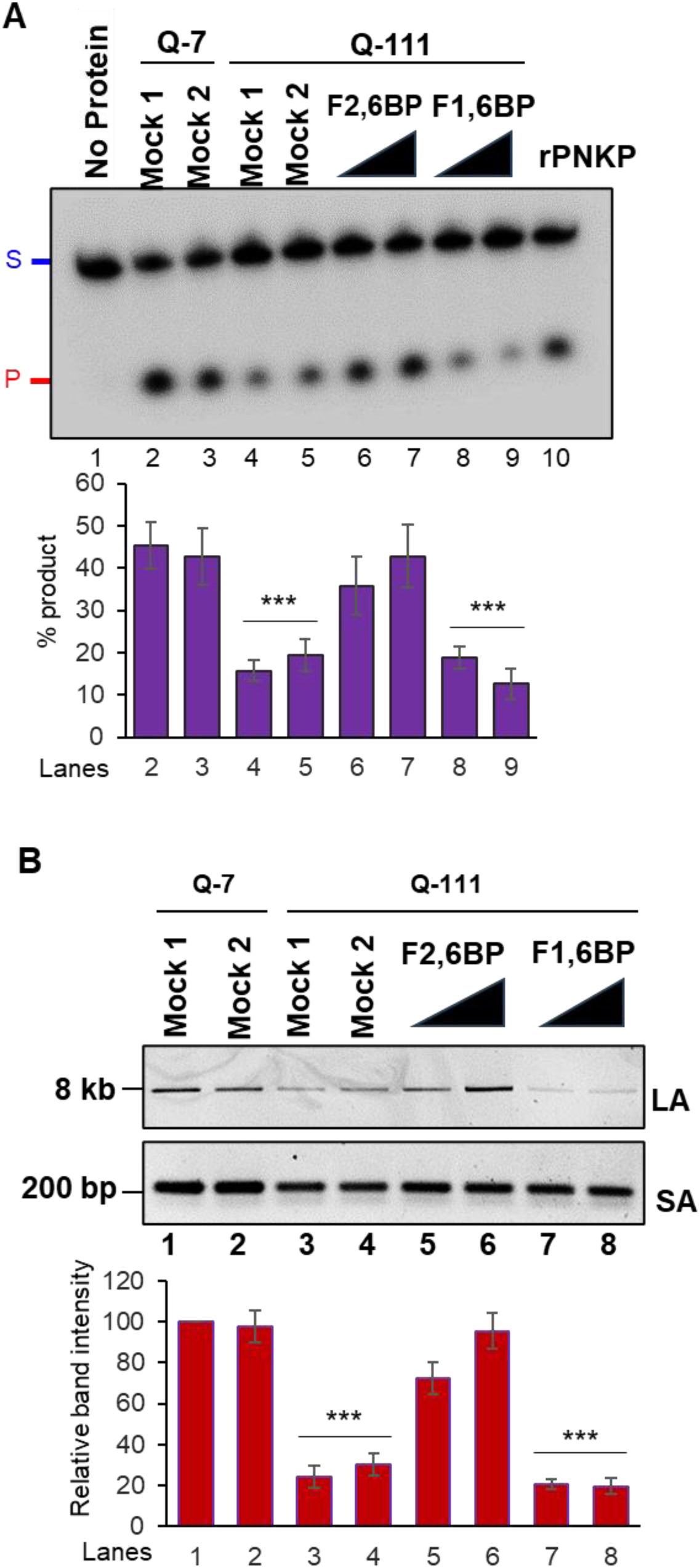
**A. Upper panel:** Representative gel image of 3’-phosphatase activity of PNKP in the mitochondrial extract of Q-7 (lanes 2-3) and Q-111 (lanes 4-9) cells with mock treatment (Mock 1; lanes 2 and 4), treatment with K16ApoE carrier peptide alone (25 µM; Mock 2; lanes 3 and 5) or supplemented with F2,6BP (100 µM; 72 and 96 h) (lanes 6-7), or F1,6BP (lanes 8-9, 100 µM) in the presence of carrier peptide. Lane 1: substrate only. Lane 10: purified PNKP (2 ng). **Lower panel:** Quantitation of the % released phosphate in the indicated lanes. Error bars show ±SD of the mean; ***P<0.005 compared to lanes 2, 3. **B.** Upper panel: Amplification of long amplicon (LA) and a short amplicon (SA) of the mitochondrial fragments to assess DNA strand break accumulation. Lower panel: The bar diagram represents the normalized (with short amplicon) relative band intensity with the mock-treated Q-7 sample arbitrarily set as 100. Error bars show ±SD of the mean; ***P<0.005 compared to lanes 1, 2.

### Exogeneous F2,6BP suppresses pathogenic HTT aggregate formation

Live cell imaging further demonstrated that the treatment with F2,6BP could partially restore the mitochondrial integrity and PFKFB3 level in Q-111 cells **(Fig. 2G, 2H)** consistent with its effect on restoration of PNKP activity and repair of mtDNA damage. Consequently, we observed a decreased association of PFKFB3 in the PNKP immunocomplex from mitochondrial extract of Q-111 cells **(Fig. 6A, lane 8 vs lane 7**); however, such association was restored in F2,6BP-treated Q-111 cells **(Fig. 6A, lane 9 vs lane 8)**. This restoration in PFKFB3 level was accompanied by a significant increase of mitochondrial membrane potential **(Fig. 3D)** and decrease in mtROS level **(Fig. 3H)** in Q-111 cells. Notably, mitochondrial respiration, as reflected in OCR, non-mitochondrial oxygen consumption, basal and maximal respiration and spare respiratory capacity showed significant enhancement upon supplementation of F2,6BP in Q-111 cells **(Fig. 3I-M)**.

**Figure 6:**
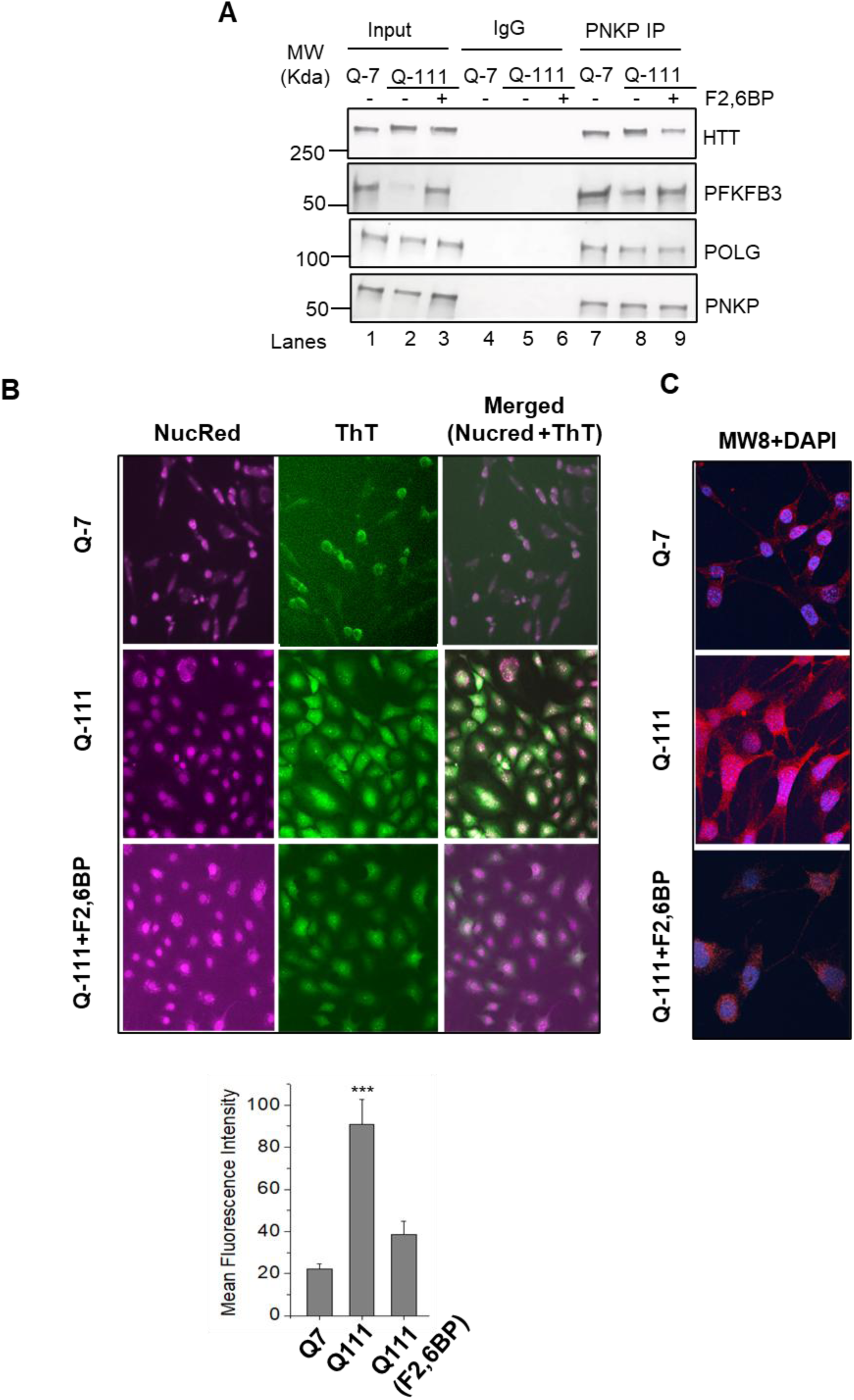
**A.** Benzonase-treated mitochondrial extracts from Q-7, mock and F2,6BP-treated (100 µM, 72 h) Q-111 cells were immunoprecipitated (IP’d) with anti-PNKP antibody (PNKP IP); or control IgG and tested for the presence of associated proteins (shown in the right) using specific Abs. **B.** HTT aggregation levels in Q-7 **(top)**, Q-111 **(middle)** and Q-111 cells treated with F2,6BP (100 µM; **bottom**) were assessed with ThT dye, which fluoresces green, and nuclear localization was indicated by NucRed dye, providing a cyan signal in the merged images. Imaging was performed on the EVOS™ M5000 system at 200X magnification. **(Bottom)** Relative quantification of the mean fluorescence intensity (in arbitrary units) (***p<0.005, between Q-7 and Q-111 cells and Q-111 cells + F2,6BP. **C.** Immunofluorescence micrographs show HTT expression and aggregation in Q-7 **(top)**, Q-111 **(middle)** and Q-111 cells treated with F2,6BP **(bottom)**. Cells were stained with Anti-HTT antibody MW8, visualized using a mouse secondary antibody conjugated to Alexa Fluor™568, resulting in red fluorescence. The nuclei were counter stained with DAPI. Images were captured at 600X magnification using a SoRa super-resolution spinning disk confocal system with motorized FRAP/photobleaching.

The polyQ expanded HTT proteins are known to be cleaved at the N-terminus, forming pathogenic aggregates, a hallmark of HD pathology^31,32^. To monitor these pathogenic protein aggregates, we used Thioflavin-T (ThT), a benzothiazole dye considered a ‘gold standard’ for detecting neurodegenerative aggregates due to its affinity for β-sheet structures. Microscopic analyses following co-staining with ThT and NucRed (a nuclear specific dye) showed significant increase in aggregate formation in Q-111 cells **(Fig. 6B, Middle Panel)** compared to Q-7 cells **(Fig. 6B, Upper Panel)**. Remarkably, F2,6BP treatment substantially reduced aggregate formation in Q-111 cells **(Fig. 6B, Lower Panel)**. We also used a monoclonal antibody (MW8) that efficiently recognizes mHTT aggregates. Staining revealed aggregate formation throughout Q-111 cells including the nuclei. The treatment of Q-111 cells with F2,6BP markedly reduced MW8 staining suggesting reduced aggregation **(Fig. 6C)**. Taken together, these results indicate a correlation between HTT aggregate formation, PFKFB3 degradation and impairment of PNKP-mediated DNA strand break repair under pathological conditions, which can be significantly rescued by F2,6BP supplementation.

### F2,6BP supplementation restores mitochondrial DNA integrity in *Drosophila* HD models

In our previous study, we demonstrated that the pan-neuronal expression of *Htt*128Q (BDSC# 33808 x 458) resulted in significant impairment of motor neuron function compared to the control used to generate the strains (*w^1118^*). When flies were supplemented with F2,6BP for 21 days, vs mock buffer control, motor function was restored, as shown by the climbing assay results^22^. To further investigate whether F2,6BP supplementation would alleviate mitochondrial repair deficiencies, we examined the DNA strand break accumulation in *Drosophila* mitochondrial genome by LA-qPCR. Flies expressing *Htt*128Q and treated with mock buffer showed elevated level of mtDNA damage **(Fig. 7, lane 2 and 4 vs lane 1),** indicating that the mock buffer could not restore mitochondrial genome integrity. However, significant repair was observed following F2,6BP supplementation and the genome integrity was comparable to the *W^1118^* flies **(Fig. 7, lane 3 and 5 vs lane 1)**. These findings confirm the ability of F2,6BP to specifically reverse the HD neurodegenerative phenotype *in vivo* in neuronal cells by restoring nuclear and mitochondrial DNA repair.

**Figure 7:**
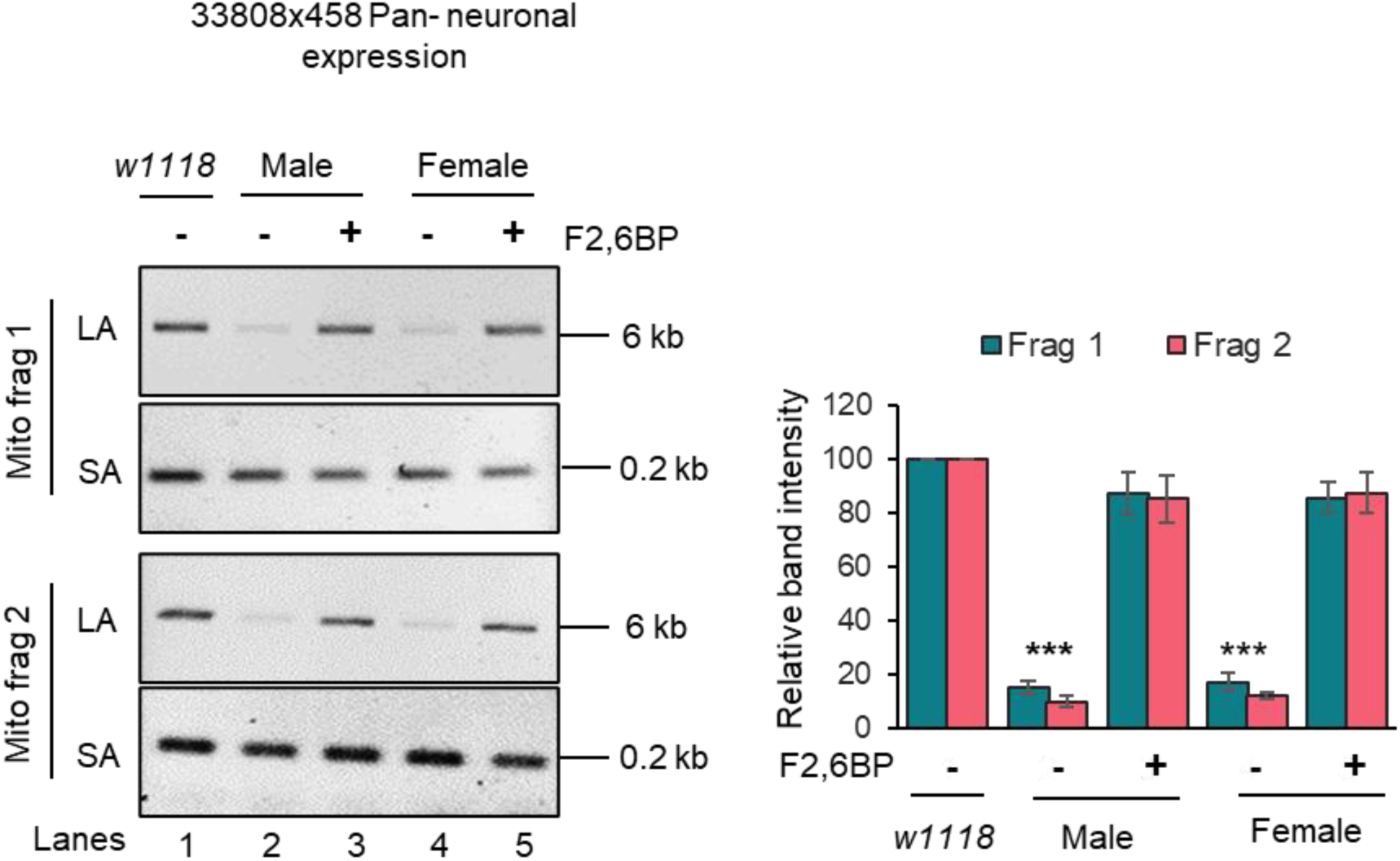
**Left panel:** Representative agarose gel images of long (LA) and short (SA) amplicon of the two mitochondrial fragments from genomic DNA of male (lanes 2-3) and female (lanes 4-5) flies with pan-neuronal expression of *Htt*128Q either mock-treated (lanes 2 and 4) or treated with F2,6BP (lanes 3 and 5). Lane 1: *w1118* (males and females). Right panel: The normalized relative band intensities were represented in the bar diagram with *w1118* arbitrarily set as 100 (error bars represent ±SD of the mean). The damage for each gene in *Drosophila* pan-neuronal expression of *Htt*128Q was significant (***P<0.005) compared to the *w1118* samples. Also, the strand breaks were significantly repaired in F2,6BP-treated samples.

Our previous study demonstrated that PFKFB3, an enzyme traditionally known for its glycolytic function, translocates to the nucleus and participates in nuclear DNA repair via transcription-coupled non-homologous end-joining (TC-NHEJ) pathways^22^. We hypothesized that locally produced F2,6BP is transferred directly to PNKP within the TC-NHEJ complex to potentiate 3’-phosphatase activity of PNKP. This process is severely impaired under HD pathogenic conditions. Our present study provides compelling evidence that PFKFB3 and its product F2,6BP are essential also for maintaining mitochondrial PNKP activity, thereby supporting efficient mitochondrial genome repair. Our findings further suggest that this repair function of PFKFB3 and F2,6BP is crucial for mitochondrial health and function. Notably, we observed that F2,6BP treatment in cells partially restores mitochondrial integrity, membrane potential and reduces the pathogenic aggregate formation in neuronal cells expressing polyQ-expanded HTT. It was generally believed that PFKFB3 levels are generally low in neurons compared to other brain cells, such as astrocytes, but now has been established that PFKFB3 is stabilized during brain excitation^33^. Our results are consistent with this report as it shows that PFKFB3’s level is low both in the nucleus and in mitochondria under pathogenic condition but not in post-mitotic healthy neurons. However, the mechanistic basis of significantly reduced PFKFB3 protein level in mitochondrial extract from HD patients requires further investigation.

Mitochondria are the sites of cellular energy production. However, recent studies provide evidence of a far wider range of mitochondrial functions. Among these, the roles of mitochondria in intracellular signaling and in communication with other cellular organelles have been elucidated in various organisms^34^. In particular, stressed mitochondria appear to induce beneficial response through effective mitochondrial-nuclear communication. This process is called “Mitochondrial retrograde signaling”- a mechanism by which mitochondria communicate to the nucleus about their functional status^35^. Our results suggest that a small molecule metabolite, F2,6BP can coordinate such communication by potentiating DNA repair in both these organelles. Currently, no curative therapy is available to halt or reverse HD pathology. Previously, a synthetic small molecule, XJB-5-131 (developed by McMurray’s group), a radical scavenger and uncoupler of oxidative phosphorylation, showed promise as a therapeutic compound by suppressing disease phenotypes in HD mouse models^36-37^. Therefore, rescuing diseased cells from oxidative DNA damage could provide a therapeutic approach. Our results in Q-111 cells and in *Drosophila* provide an avenue of rescue of oxidative stress-induced nuclear and mitochondrial DNA damage through F2,6BP supplementation, concurrent with improved mitochondrial functionality. Importantly, an endogenous, naturally occurring molecule produced in the body can essentially harness the body’s own mechanisms to promote healing. This “therapeutics from within” approach can involve manipulating the levels of F2,6BP to achieve therapeutic benefit. Thus, our results lay the groundwork for exploring a promising therapeutic approach through F2,6BP supplementation or a non-toxic analog to prevent key aspects of HD progression by maintaining nuclear and mitochondrial genomic integrity.

Recent reports have demonstrated that medium spiny neurons (MSNs) are the most vulnerable cell types in HD striatum^38-39^. Cholinergic neurons, despite somatic expansion of CAG repeats, are not significantly lost in HD striatum. The authors concluded that somatic repeat expansion is required but not sufficient for neuronal loss. We postulate that differential sensitivity of various cell types is attributed to the residual F2,6BP level and consequent PNKP activity in both mitochondria and nucleus. We predict that degradation of PFKFB3 and the resultant decrease in intracellular F2,6BP level in MSNs is severe, affecting PNKP activity leading to persistent DSBs and cell death. Measuring F2,6BP level and PNKP activity in both the organelles of those vulnerable cells warrants further investigation.

## Methods

### Cell culture

Human Embryonic Kidney 293 (HEK293; ATCC CRL-1573) cells were grown at 37°C and 5% CO_2_ in DMEM: F-12 (1:1, Cellgro) medium containing 10% fetal bovine serum (Sigma), 100 units/ml penicillin, and 100 units/ml streptomycin (Cellgro). Mouse striatum derived cell line from a knock-in transgenic mouse containing homozygous Huntingtin (HTT) loci with a humanized Exon 1 containing 7 or 111 polyglutamine repeats (Q-7 and Q-111; Corriell Institute; Cat# CH00097 and CH00095, respectively) were cultured and maintained in Dulbecco Modified Eagles Medium (high glucose) with 2 mM L-glutamine containing 10% fetal bovine serum, 100 units/ml penicillin and streptomycin, and 0.4 mg/ml G418. Q-111 cells lose the ability to proliferate and survive. Therefore, high-passage numbers were avoided. We routinely tested for mycoplasma contamination in cultured cells using the Mycoalert Mycoplasma Detection Kit (Lonza) according to the manufacturer’s protocol, and the cells were found to be free from mycoplasma contamination.

### Isolation of mitochondria and mitochondrial extracts from cultured cells and post-mortem human tissues

Mitochondria were extracted from cultured cells and post-mortem tissues using mitochondria isolation kits from Thermo Fischer (Cat# 89874 and 89801). Isolated mitochondria were washed with PBS, treated with trypsin (1 mg/ml in PBS) for 15 min at room temperature to remove contaminating proteins adhering to the outer surface of mitochondria, and then extensively washed with PBS. The washed mitochondria were lysed in 50 mM Tris, pH 7.5, 150 mM NaCl, 1 mM EDTA, 1 mM DTT, and 1% Triton X-100 to prepare purified mitochondrial extract.

### Immunoblotting

The proteins in the mitochondrial extracts were separated onto a Bio-Rad 4-20% gradient Bis-Tris gel, then electro-transferred on a nitrocellulose (0.45 μm pore size; GE Healthcare) membrane using 1X Bio-Rad transfer buffer. The membranes were blocked with 5% w/v skimmed milk in TBST buffer (1X Tris-Buffered Saline, 0.1% Tween 20), then immunoblotted with appropriate antibodies [PNKP (BB-AB0105, BioBharati Life Science), PFKFB3 (GTX108335, GeneTex), COX4 (GTX114330, GeneTex)]. The membranes were extensively washed with 1% TBST followed by incubation with anti-isotype secondary antibody (GE Healthcare) conjugated with horseradish peroxidase in 5% skimmed milk at room temperature. Subsequently, the membranes were further washed three times (10 min each) in 1% TBST, developed using ECL^TM^ Western Blotting Detection Reagents (RPN2209, GE Healthcare) and imaged using Kwikquant image analyzer and image analysis software (ver 5.2) (Kindle Biosciences). Cytosolic and nuclear extracts were prepared as described earlier^22^ and their purity was checked by immunoblotting with GAPDH (BB-AB0060 BioBharati Life Science) and HDAC2 (GTX109642, GeneTex) antibodies, respectively.

### Immunopulldown (IP)

Mitochondrial extracts from Q-7 and Q-111 cells were IP’d using PFKFB3 and PNKP Abs and with Protein A/G PLUS agarose beads (Sc-2003, Santa Cruz Biotechnology) overnight, followed by four washes with Wash buffer [20 mM HEPES (pH 7.9), 150 mM KCl, 0.5 mM EDTA, 10% glycerol, 0.25% Triton-X-100 and 1X protease inhibitors] and eluted with Laemmli Sample Buffer (Bio Rad; final concentration 1X). The immunoprecipitates were tested for the interacting proteins using appropriate Abs [PFKFB3, HTT, PNKP, POLG (MA5-57385), Lig IIIα (in-house) and RNAPII (920202, pSer2, H5 Ab).

### Human tissue samples

Deidentified human post-mortem frontal cortex tissue of HD patients (detailed in our earlier publication)^22^ and age-matched controls (IRB exempt) were obtained from the biorepository of the Michigan Brain Bank, USA through a Materials Transfer Agreement (UTMB 22-UFA00474).

### Assay of 3’-phosphatase of mtPNKP

The 3’-phosphatase activity of PNKP in the mitochondrial extract of post-mortem patients’ frontal cortex and age-matched control subjects or Q-7 or Q-111 cells (250 ng) or with purified recombinant PNKP (2 ng) was conducted as we described previously^22^. Five pmol of the radiolabeled substrate was incubated at 37°C for 15 min in buffer A (25 mM Tris-HCl, pH 8.0, 100 mM NaCl, 5 mM MgCl_2_, 1 mM DTT, 10% glycerol and 0.1 μg/μl acetylated BSA). 5 pmol of non-radiolabeled substrate was used as cold substrate. For *in vitro* PNKP restoration, similar assays were done after incubation of F2,6BP/F6P/F1,6BP (in amounts as indicated in the figure legends) with the mitochondrial extracts for 15 min. The radioactive bands were visualized using a PhosphorImager (GE Healthcare) and quantitated using ImageQuant software. The data are represented as % product (released phosphate or kinase product) released from the radiolabeled substrate with a value arbitrarily set at 100%.

### Enzymatic preparation of F2,6BP

Enzymatic preparations of F2,6BP were conducted following the protocol as described before^22^.

### Estimation of F2,6BP in patients’ extracts

The quantitation of F2,6BP from the purified mitochondrial extract of post-mortem patients was performed as described before^22^.

### Long amplicon quantitative PCR (LA-qPCR)

Genomic DNA was extracted from post-mortem tissues, striatal neuronal cells and *Drosophila* (10 each) using the Genomic tip 20/G kit (Qiagen) per the manufacturer’s protocol, to ensure minimal DNA oxidation during the isolation steps. The DNA was quantitated by Pico Green (Molecular Probes) in a black-bottomed 96-well plate and gene-specific LA qPCR assays were performed as described earlier^22^ using Long Amp Taq DNA Polymerase (New England BioLabs). The amplified products were then visualized on gels and quantitated with ImageJ software (NIH). The extent of damage was calculated in terms of relative band intensity with a control/mock-treated sample or *w^1118^* sample (for *Drosophila* studies) considered as 100. All oligos (LA: long amplicon and SA: short amplicon) used in this study are listed below:

**Oligo sets for amplification of mitochondrial fragment from mouse genome (Q-7 and Q-111 cells):** For LA PCR: LA 1: 5′-GCC AGC CTG ACC CAT AGC CAT ATT AT-3′ LA 2: 5′-GAG AGA TTT TAT GGG TGT ATT GCG G-3′ For SA PCR: SA 1: -CCC AGC TAC TAC CAT CAT TCA AGT-3′ SA 2: 5′-GAT GGT TTG GGA GAT TGG TTG ATG-3′

**Oligo sets for amplification of mitochondrial fragment from human genome (post-mortem patients):** For LA PCR: LA 1: 5’-TCTAAGCCTCCTTATTCGAGCCGA-3’ LA 2: 5’-TTTCATCATGCGGAGATGTTGGATGG-3’

For SA PCR: SA 1: 5’-CCCCACAAACCCCATTACTAAACCCA-3’ SA 2: 5’-TTTCATCATGCGGAGATGTTGGATGG-3’.

**Oligo sets for amplification of mitochondrial fragment from *Drosophila* genome:** For Fragment 1: LA 1: 5’-TGTGAATAATAGCCCCAGCACA-3’ LA 2: 5’-GCTGGAATGAATGGTTGGACG-3’ SA 1: 5’-ACACCTGCCCATATTCAACCA-3’ SA 2: 5’-ACTGGTCGAGCTCCAATTCA-3’; For Fragment 2: LA 3: 5’-GTGAATAATAGCCCCAGCACA-3’ LA 4: 5’-AGGCTGGAATGAATGGTTGGA-3’ SA 3: 5’-ACCTGCCCATATTCAACCAGA-3’ SA 4: 5’-TCAACTGGTCGAGCTCCAAT-3’.

### Binding of F2,6BP to PNKP

Addition of F2,6BP (400 nM) to PNKP resulted in partial quenching of the tryptophan fluorescence of PNKP at 340 nm, when excited at 295 nm with no change in the emission maximum. Binding affinity (K_d_) was determined by following fluorescence quenching (a measure of ligand binding) as a function of F2,6BP concentration. The maximum quenching of fluorescence intensity observed at saturating concentration of F2,6BP was taken as 1, and the observed quenching at different concentrations of F2,6BP was plotted as the fraction bound *versus* free F2,6BP concentration. Binding of F2,6BP to PNKP was analysed using GraphPad Prism Software, San Diego, CA using the equation:

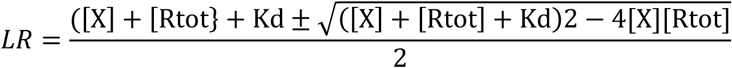

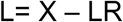

where X is the total ligand concentration and Rtot is the total receptor concentration (Same units as X). LR refers to ligand bound to the receptor and L is the free ligand concentration.

### *In cell* delivery of the exogenous F2,6BP/F1,6BP in Q-7 and Q-111 cells

For delivery of the glycolytic metabolites into the mouse striatum-derived neuronal cells (Q-7 and Q-111), we followed the protocol optimized in our lab^20^. Briefly, 200 μM of F2,6BP or F1,6BP was mixed with 25 μM cell-permeable carrier peptide K16ApoE (Mayo Proteomic Core Facility) and incubated for 45 min at RT; then metabolite-peptide mix was added to the cells and incubated for 6 h in reduced serum (oPTIMEM) media. After 6 h, FBS containing complete media was added and cells were harvested at 72 and 96 h for DNA repair assays/LA-qPCR.

### Antibodies and chemicals for immunostaining and flow cytometry

The antibodies and chemicals used for immunostaining were as follows: Mouse Anti-HTT antibody-MW8 (Creative Biolab, San Diego, USA), Tetramethylrhodamine Methyl Ester Perchlorate (TMRM) from ThermoFischer scientific, USA, Thioflavin-T (Sigma-Aldrich, USA), DAPI (Novus, USA), NucRed dye (647) and MitoTracker from Invitrogem, USA).

### FACS analysis

TMRM is widely used for assessing mitochondrial membrane potential due to its ability to accumulate within mitochondria in proportion to membrane potential, as predicted by the Nernst equation. The cells were grown on 10 cm cell culture dishes in DMEM at 37°C with 5% CO₂. The staining protocol was as follows: Briefly, TMRM (1 μM) stock solution and Verapamil (2 nM) were mixed in a ratio of 2:1 and the cells were incubated in the staining solution for 1 hour at 37°C. Fluorescence intensity was measured by flow cytometry (FACSCalibur, BD Biosciences, CA, USA) using CellQuest Pro software (version 5.2, BD Biosciences, Franklin Lakes, NJ, USA). Each experiment was performed in duplicate.

### Estimation of Mitochondrial ROS level

Reactive oxygen species (ROS) levels in tissue/cell extracts were estimated using the fluorescent dye H₂DCFDA. Tissue samples were homogenized in ice-cold phosphate-buffered saline (PBS), centrifuged at 10,000 × g for 10 minutes at 4°C, and the supernatant was collected. The extracts were incubated with 10 µM H₂DCFDA at 37°C for 30 minutes in the dark. Fluorescence intensity was measured using a microplate reader at 485 nm excitation and 530 nm emission wavelengths. ROS levels were expressed as relative fluorescence units (RFU) normalized to total protein content. Data represents the mean of 14 diseased and 14 control samples, each analyzed in triplicate, and are presented with standard error.

### Mitochondrial respiration by sea-horse analyses

Mitochondrial respiratory function was evaluated in Q-7 (wild-type) and Q-111 (HD) cell lines using the Seahorse XF96 Cell Mito Stress Test. Q-111 cells were treated with F2,6BP (200 µM) and allowed to recover for 72 hours prior to analysis. Both untreated Q-7 and Q-111 cells, as well as F2,6BP-treated Q-111 cells, were plated at a density of 25,000 cells per well in Seahorse XF96 plates. Oxygen consumption rate (OCR) was measured under basal conditions and in response to sequential injections of mitochondrial modulators. First, oligomycin was added to inhibit ATP synthase, allowing for assessment of ATP-linked respiration through the corresponding drop in OCR. This was followed by the injection of FCCP, a mitochondrial uncoupler that induces maximal respiration by collapsing the proton gradient, revealing the spare respiratory capacity. Finally, antimycin A, a complex III inhibitor, was introduced to inhibit the electron transport chain, enabling the determination of non-mitochondrial respiration by analyzing the differences in OCR between oligomycin and antimycin A treatments. This approach allowed for comparative evaluation of basal respiration, maximal respiratory capacity, and mitochondrial efficiency between the two cell lines and treatment conditions.

## Author contribution statement

TH, GG and AC conceived the research. TH, GG, AC, SM, MLH, MW and BK designed the research. AC, SM, MM, RSM, SS, TB, MK, SGS, NT performed research. AC, SM, NT, MW, GG and TH wrote the manuscript. All the authors read and approved the final version of the manuscript.

## Data availability

All study data are included in the article.

## Acknowledgement

This work was supported by National Institute of Health Grants 2R01 NS073976 and R56NS073976 to TH, R01AI163327 and R21 AG078635 to GG, Don and Nancy Mafrige Professorship in Neurodegenerative Diseases (BK), Alzheimer’s Association Research Grant (AARG-17-533363 BK), National Institutes of Aging (R21-AG059223 and R01-AG063945 BK), National Institute of Health grant RF1NS112719 to MLH, Canadian Institutes of Health Research grant PJT168869 and University of Alberta bridge grant ZAMPD to MW. We thank the Michigan Brain Bank (5P30 AG053760 University of Michigan Alzheimer’s Disease Core Center) for providing us with the tissue of post-mortem HD patients and their age-matched controls.

## References

1. H. Y. Zoghbi, H. T. Orr, Glutamine repeats and neurodegeneration. Annu Rev Neurosci. 23:217–47 (2000).

2. Genetic Modifiers of Huntington’s Disease, C. Identification of Genetic Factors that Modify Clinical Onset of Huntington’s Disease. Cell 162, 516-526, doi: 10.1016/j.cell.2015.07.003 (2015).

3. Genetic Modifiers of Huntington’s Disease Consortium. Electronic address, g. h. m. h. e. & Genetic Modifiers of Huntington’s Disease, C. CAG Repeat Not Polyglutamine Length Determines Timing of Huntington’s Disease Onset. Cell 178, 887-900 e814, doi: 10.1016/j.cell.2019.06.036 (2019).

4. R. A. Quintanilla, G. V. W. Johnson, Role of mitochondrial dysfunction in the pathogenesis of Huntington’s disease. Brain Res Bull Oct 28;80(4-5):242–7 (2009).

5. A. Jurcau, C. M. Jurcau, Mitochondria in Huntington’s disease: implications in pathogenesis and mitochondrial-targeted therapeutic strategies. Neural Regen Res Jul;18(7):1472–1477 (2023).

6. A. Neueder et al. Huntington’s disease affects mitochondrial network dynamics predisposing to pathogenic mitochondrial DNA mutations. Brain Jun 3;147(6):2009–2022 (2024).

7. E. Martin-Solana et al. Disruption of the mitochondrial network in a mouse model of Huntington’s disease visualized by in-tissue multiscale 3D electron microscopy. Acta Neuropathol Commun Jun 5;12(1):88 (2024).

8. Y. Dai et al. A comprehensive perspective of Huntington’s disease and mitochondrial dysfunction. Mitochondrion May 70:8–19 (2023).

9. S. Ayala-Peña, Role of oxidative DNA damage in mitochondrial dysfunction and Huntington’s disease pathogenesis. Free Radic Biol Med Sep:62:102–110 (2013).

10. I. Shokolenko et al. Oxidative stress induces degradation of mitochondrial DNA. Nucleic Acids Res May;37(8):2539–48 (2009).

11. K. Acevedo-Torres et al. Mitochondrial DNA damage is a hallmark of chemically induced and the R6/2 transgenic model of Huntington’s disease. DNA Repair (Amst) Jan 1;8(1):126–36 (2009).

12. A. Siddiqui et al. Mitochondrial DNA damage is associated with reduced mitochondrial bioenergetics in Huntington’s disease. Free Radic Biol Med. Oct 1;53(7):1478–88 (2012).

13. L. Weissman et al. DNA repair, mitochondria, and neurodegeneration. Neuroscience Apr 14;145(4):1318–29 (2007).

14. A. Jilani, et al., Molecular cloning of the human gene, PNKP, encoding a polynucleotide kinase 3’-phosphatase and evidence for its role in repair of DNA strand breaks caused by oxidative damage. J Biol Chem Aug 20;274(34):24176-86 (1999).

15. F. Karimi-Busheri, et al., Molecular characterization of a human DNA kinase. J Biol Chem Aug 20;274(34):24187–94 (1999).

16. M. Shimada, L. C. Dumitrache, H. R. Russell, P. J. McKinnon, Polynucleotide kinase-phosphatase enables neurogenesis via multiple DNA repair pathways to maintain genome stability. Embo Journal 34, 2465–2480, doi:10.15252/embj.201591363 (2015).

17. J. Shen, et al., Mutations in PNKP cause microcephaly, seizures and defects in DNA repair. Nat Genet 42, 245–249 (2010).

18. C. Poulton, et al., Progressive cerebellar atrophy and polyneuropathy: expanding the spectrum of PNKP mutations. Neurogenetics 14, 43–51 (2013).

19. A. Islam, et al., Functional analysis of a conserved site mutation in the DNA end processing enzyme PNKP leading to ataxia with oculomotor apraxia type 4 in humans. J Biol Chem 299, 104714 (2023).

20. B. Jiang et al. Mutations of the DNA repair gene PNKP in a patient with microcephaly, seizures, and developmental delay (MCSZ) presenting with a high-grade brain tumor. Sci Rep. Mar 30;12(1):5386 (2022).

21. R. Gao, et al., Mutant huntingtin impairs PNKP and ATXN3, disrupting DNA repair and transcription. Elife 8, doi:10.7554/eLife.42988 (2019).

22. A. Chakraborty et al. Fructose-2,6-bisphosphate restores DNA repair activity of PNKP and ameliorates neurodegenerative symptoms in Huntington’s disease. Proc Natl Acad Sci USA Sep 24;121(39):e2406308121 (2024).

23. J. Hu et al. Repair of formamidopyrimidines in DNA involves different glycosylases: role of the OGG1, NTH1, and NEIL1 enzymes. J Biol Chem Dec 9;280(49):40544-51 (2005).

24. S. M. Mandal et al. Role of human DNA glycosylase Nei-like 2 (NEIL2) and single strand break repair protein polynucleotide kinase 3’-phosphatase in maintenance of mitochondrial genome. J Biol Chem. Jan 20;287(4):2819–29 (2012).

25. N. Tahbaz et al. Role of polynucleotide kinase/phosphatase in mitochondrial DNA repair. Nucleic Acids Res Apr;40(8):3484–95 (2012).

26. A.J. Neil et al. Transcription blockage by bulky end termini at single-strand breaks in the DNA template: differential effects of 5’ and 3’ adducts. Biochemistry Nov 6;51(44):8964–70 (2012).

27. A. Chakraborty, et al., Classical non-homologous end-joining pathway utilizes nascent RNA for error-free double-strand break repair of transcribed genes Nature communications 7, 13049, doi:10.1038/ncomms13049 (2016).

28. S.L. Chalasani et al. Persistent 3’-phosphate termini and increased cytotoxicity of radiomimetic DNA double-strand breaks in cells lacking polynucleotide kinase/phosphatase despite presence of an alternative 3’-phosphatase. DNA Repair (Amst). Aug;68:12–24 (2018).

29. E. Romero, et al., Suppression of neurodegeneration and increased neurotransmission caused by expanded full-length huntingtin accumulating in the cytoplasm. Neuron Jan 10;57(1):27–40 (2008).

30. F. Trettel, et al., Dominant phenotypes produced by the HD mutation in STHdh(Q111) striatal cells. Human molecular genetics 9:2799–809 (2000).

31. M. Z. Chen et al. N-Terminal Fragments of Huntingtin Longer than Residue 170 form Visible Aggregates Independently to Polyglutamine Expansion. J Huntingtons Dis; 6(1):79–91 (2017).

32. S. J. Tabrizi et al. Huntington disease: new insights into molecular pathogenesis and therapeutic opportunities. Nat Rev Neurol Oct;16(10):529–546 (2020).

33. C. M. Diaz-Garcia, et al., Neuronal Stimulation Triggers Neuronal Glycolysis and Not Lactate Uptake. Cell Metab 26, 361–374 e364, doi: 10.1016/j.cmet.2017.06.021 (2017).

34. K. H. Kim, C. B. Lee, Socialized mitochondria: mitonuclear crosstalk in stress. Exp Mol Med May;56(5):1033–1042 (2024).

35. R. Desai, et al. Mitochondria form contact sites with the nucleus to couple prosurvival retrograde response. Sci Adv Dec 18;6(51): eabc9955 (2020).

36. Z. Xun, et al., Targeting of XJB-5-131 to mitochondria suppresses oxidative DNA damage and motor decline in a mouse model of Huntington’s disease. Cell Rep Nov 29;2(5):1137–42 (2012).

37. P. Wipf, et al., A Double-Pronged Sword: XJB-5-131 Is a Suppressor of Somatic Instability and Toxicity in Huntington’s Disease. J Huntingtons Dis 11(1):3–15 (2022).

38. A. Jiang, et al., Single nuclei RNA-seq reveals a medium spiny neuron glutamate excitotoxicity signature prior to the onset of neuronal death in an ovine Huntington’s disease model. Hum Mol Genet Aug 18;33(17):1524–1539 (2024).

39. K. Mätlik, et al., Cell-type-specific CAG repeat expansions and toxicity of mutant Huntingtin in human striatum and cerebellum. Nat Genet Mar;56(3):383–394 (2024).

